# Modeling human echolocation using a Kalman filter

**DOI:** 10.64898/2026.07.01.735693

**Authors:** Sofia Krasovskaya, James M. Coughlan, Santani Teng

## Abstract

Some blind individuals use echolocation, a skill that allows them to better navigate their environment using echoes from self-generated mouth clicks reflected off surrounding surfaces. Echolocation involves a complex interplay of sensory accumulation, information processing, dynamic prediction, motor planning and execution in real-time. Computational modeling offers a valuable approach to understanding the cognitive and neural mechanisms underlying echolocation performance, in particular the temporal dynamics of the process. We present a computational model of human echolocation behavior based on a Kalman filter, where we treat the echolocator as an active sensor that maintains an internal belief about the target’s location and continuously refines it via echo feedback. The model, based on observations of echolocation in blind human experts, simulates the use of mouth clicks and returning echoes to localize and orient toward a target under varying conditions. In the experiment, the target is placed at a random azimuth in the frontal plane. An echolocator aims a series of mouth clicks in various directions and infers the target azimuth using acoustic information received from the click echoes. The system integrates three major components: (1) a simulation of echoacoustic interaural time differences (ITD) to estimate the relative head-target angle; (2) a Kalman filter that processes these ITDs to iteratively update probabilistic beliefs about target location and associated uncertainty; and (3) a motor control system that modulates head movements with the current belief state. The Kalman filter serves as a representation of the internal state of the observer, where its beliefs drive the direction of head rotation, and its uncertainty estimates drive head velocity adjustments. Model performance demonstrates that simple predictive computational approaches can reproduce key aspects of echo-guided sensorimotor learning, providing a framework that may be leveraged to develop biologically plausible models, advance understanding of best practices, and potentially improve intervention strategies.

## Introduction

Blind individuals can navigate complex environments using echolocation - a skill for which one uses mouth clicks and processes echoes reflected off surrounding surfaces (Thaler & Goodale, 2016). Echolocation in real-world scenarios exemplifies *active sensing:* the brain dynamically integrates emitted clicks, noisy echoacoustic measurements, and self-motion over time to meet behavioral goals (Flanagin et al., 2017). This poses a computational challenge: how does the brain maintain and update spatial estimates from brief, noisy echoes?

Some behavioral studies have characterized human echolocation in terms of spatial performance (Kolarik et al., 2014; Teng et al., 2012), and existing computational models of echolocation have focused on aspects of the echolocation process taken in isolation, such as vocalization acoustics (Thaler et al., 2017), theoretical neuronal mechanisms (Hoshino & Kuroiwa, 2002), or psychophysical performance on specific tasks (Teng & Whitney, 2011; Teng et al., 2012; Thaler et al., 2011). Some works (García-Lázaro & Teng, 2026; Thaler et al., 2018) have suggested that perceptual evidence accumulates over multiple clicks during target localization or detection tasks. Rosenblum (2000) reported a perceptual advantage for mobile vs. stationary echolocators in an obstacle perception task. Some mobility-echolocation interactions have been described in other species, such as bats (Moss & Surlykke, 2010; Ulanovsky & Moss, 2008), and in robotic ‘terrestrial bat’ simulations maneuvering on the ground (Eliakim et al., 2018; Steckel & Peremans, 2013). Still, to our knowledge, there has been no computational framework of active sensing dynamics in echolocating humans that would capture the underlying inference and control processes. Understanding the computational principles underlying echolocation could reveal general mechanisms of active sensing, potentially shared with other modalities.

To close this gap, we adapted a modeling framework well established for sequential eye movements in vision (Mohsenzadeh et al., 2016; Renninger et al., 2007; Yang et al., 2016): sensory samples and motor movements alternating in service of a simple, relatively ecological behavioral goal. Our computational model of echolocation couples a Kalman Filter (KF) (Kalman, 1960; Welch, Bishop, et al., 1995) for probabilistic state estimation with a motor controller for goal-directed head movements (Shadmehr & Mussa-Ivaldi, 2012; Shadmehr et al., 2010; Wolpert & Kawato, 1998; Wolpert & Ghahramani, 2000). The KF framework is appropriate because it captures the process of integration of uncertain information over time, while the coupled controller architecture was inspired by theoretical proposals for sensorimotor integration (Wolpert & Kawato, 1998). The head movements generated by the model are driven by uncertainty, with larger sweeps reflecting higher uncertainty and smaller head adjustments as confidence grows. This creates a closed sensorimotor loop driven by the observer’s internal belief about the target position, which both guides and is guided by head movements and click-echo samples.

We applied this model to a previous behavioral study (Patel et al., 2024; Teng & Fusco, 2019; Teng et al., 2026), in which a blind expert observer used echolocation to orient toward a target in the frontal hemifield. Across three difficulty conditions mirroring those of the behavioral study, the model reproduced key features of expert human behavior, including directed convergence onto a target and performance degradation with decreased target dimensions. These findings demonstrate that a straightforward KF architecture coupled with uncertainty-driven motor control may provide a computationally plausible account of active sensing behavior in humans.

## Model Architecture

### Overview and Design

The model represents a simulation of participant behavior during a previously conducted echolocation experiment (Teng & Fusco, 2019; Teng et al., 2026). There, the participant was seated in a sound-insulated, echo-damped chamber. An acoustically reflective target (a board covered with aluminum foil) was placed at a random azimuth in the frontal hemifield at a distance of 1 m. The participant located the target using mouth clicks and indicated its position via head rotation and a button press to indicate the end of the trial (Fig. 1).

**Fig. 1.**
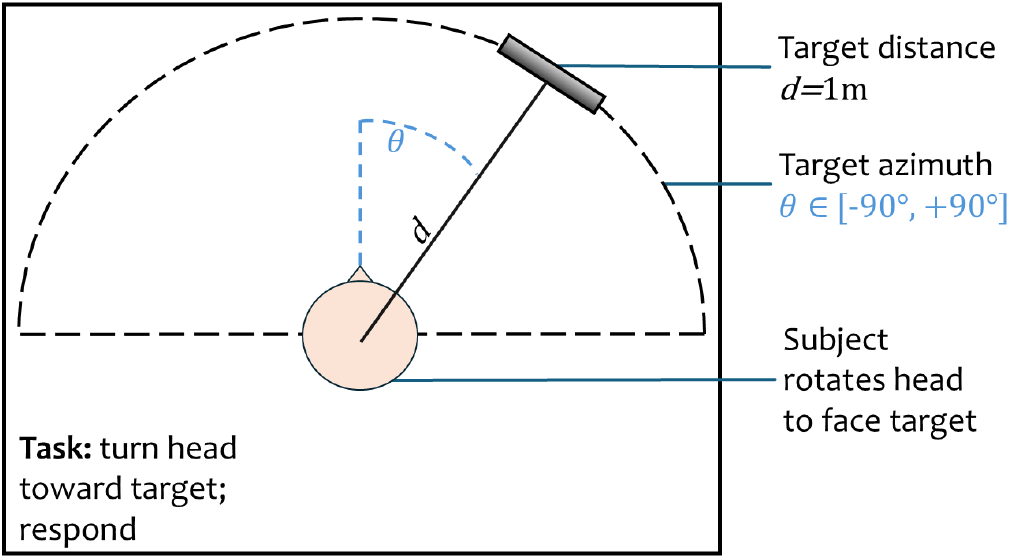
Schematic representation of the original behavioral experiment with a human participant.

The behavioral experiment included three conditions: Big Target (∼1044 cm^2^), Small Target (∼ 42.5 cm^2^) and a No-Click (control) condition, where a big target was present, but the participant could not use tongue clicks to localize the target. Mirroring this behavioral paradigm, here we evaluate model performance on Big and Small Target analogs differing only by target size (and thus the strength of the echo), and a control condition without clicks.

In the model, we treat the echolocator as an active sensor that maintains an internal belief about the location of the target and continuously refines it via echo feedback from emitted echolocation clicks. The model consists of two interacting systems: a motor system and a belief system (Fig. 2). The simulated observer’s search patterns are centered on the belief system’s current estimate 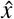, which updates with each KF measurement *z*, creating a direct coupling between perception and action as the center of motion shifts toward the true target location.

**Fig. 2.**
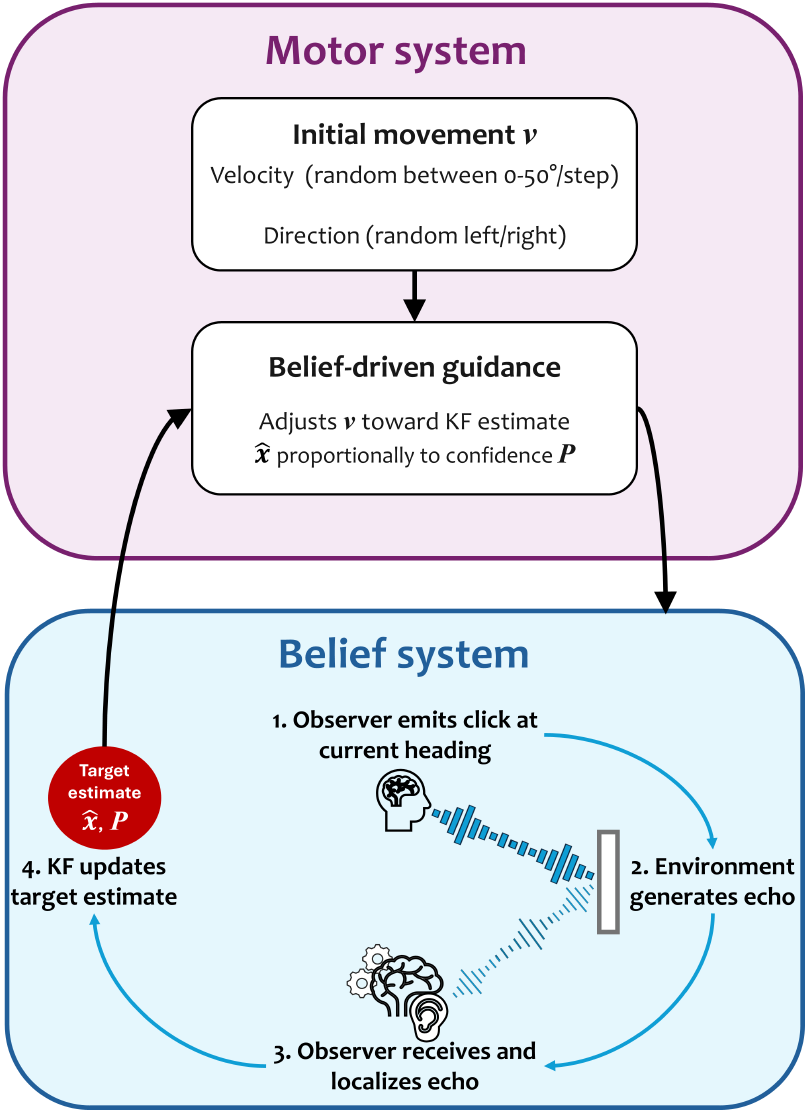
Conceptual overview of the model. The movement module (top, purple) comprises *Initial baseline movement* (*v*) modified by *Belief-driven guidance*. KF prediction and updating (bottom, blue) is triggered with each click, leading to the analysis of the resulting echo by the Acoustic measurement and processing system (*bottom, 1-3*). The direction and quality of the heading vector update the KF’s estimated target location, which in turn updates the movement program.

The model consists of two parallel systems interacting as shown in Figure 3: a motor system (left, purple) that controls head movements and a belief system (right, blue) that maintains probabilistic estimates of target location using a KF. At each timestep, the motor system operates in absolute spatial coordinates, tracking the current head azimuth θ and the desired head azimuth θ_*target*_. It reads the current target estimate 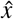 and uncertainty *P* from the KF, calculates position error ε based on the difference between θ and θ_*target*_ and updates velocity via proportional gain *K*_*p*_ and damping. Meanwhile, the belief system performs a prediction step, determines whether to emit a click based on timing parameters, and, if a click occurs, simulates an ITD-based measurement at the current head azimuth to update 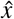 and *P*. The updated belief state (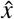, *P*) feeds back to the motor system at the next timestep. Thus, the model implements a closed-loop sensorimotor architecture that transforms acoustic input into goaldirected head movements.

**Fig. 3.**
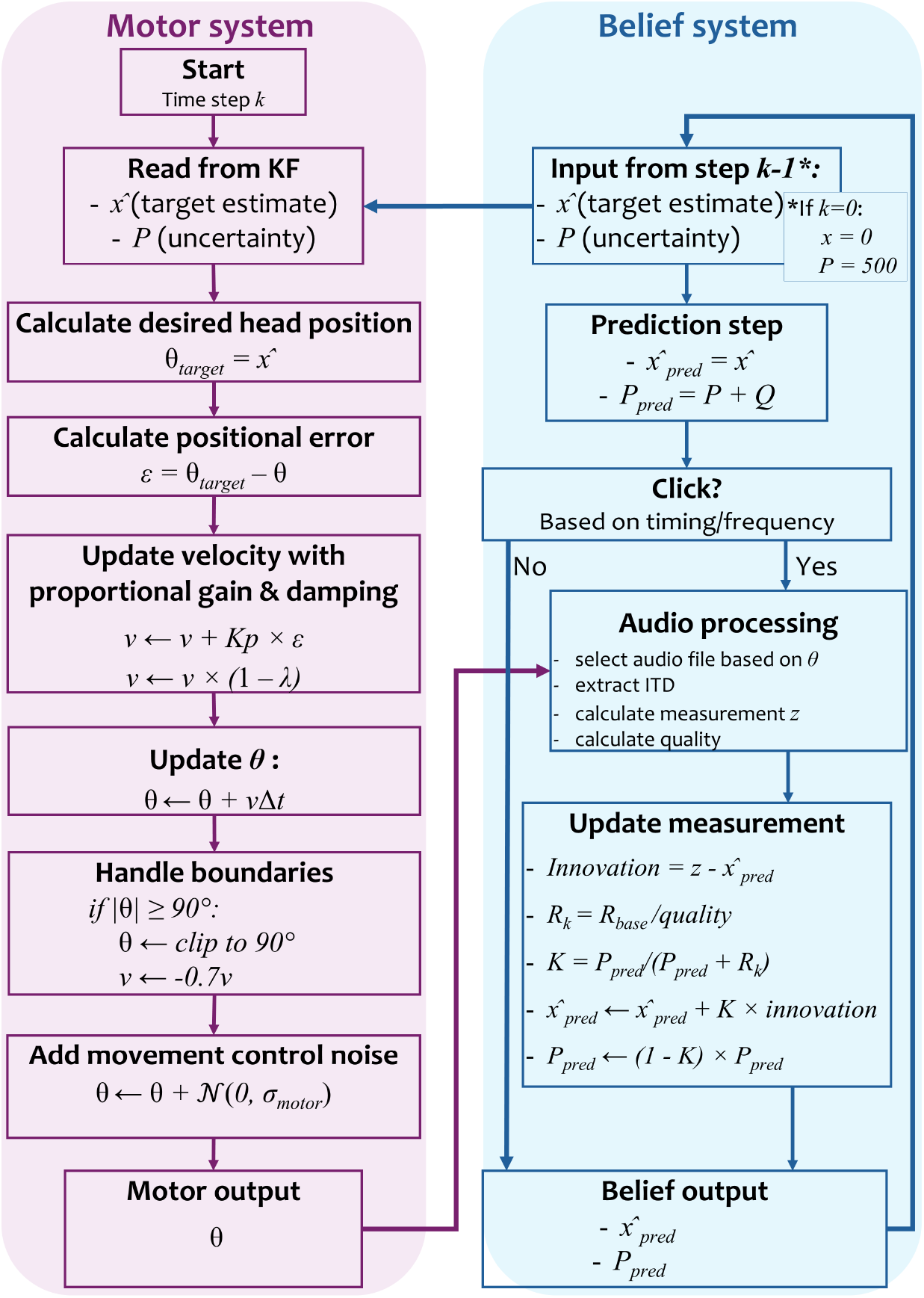
Update cycle at a single timestep *k*. At each timestep *k*, the motor system (purple) reads target estimate *x* from the KF, calculates position error, and updates head position θ through proportional velocity control with damping and boundary constraints. The belief system (blue) performs KF prediction, determines click timing and processes audio at the current head-target angle to extract ITD-based measurements that update target estimate and uncertainty. The updated belief state feeds back to the motor system, creating a closed sensorimotor loop. KF: Kalman Filter; ITD: Interaural Time Difference.

These two systems are architecturally implemented through three interconnected modules: (1) an Acoustic measurement and processing module triggered when a click is made; (2) a Belief module that manages click timing and updates the belief state based on the processed acoustic measurements; and (3) a Motor control module that moves the head based on the belief state updates. Detailed descriptions of each module are provided in their respective sections below.

### Model Parameters

To ensure consistency, we run the three conditions on one set of pregenerated parameters and target locations. The model is constrained to a set of fixed parameters, listed in Table 1. The parameters were determined through a combination of empirical constraints, theoretical considerations and iterative tuning to produce biologically realistic behavior. The model was not formally fitted to human behavioral data. Instead, parameters were adjusted iteratively to ensure that convergence behavior qualitatively matched human echolocation, response times fell within biologically plausible ranges, and target size effects emerged naturally from reflectivity scaling without condition-specific parameter changes.

**Table 1.** Model parameters.

|  | Parameter | Value | Description |
| --- | --- | --- | --- |
| Empirically constrained | $\Delta t$ | 0.1 s | Timestep. Temporal resolution of simulation balancing computational efficiency with simulation of continuous head movements and click timing. Converts between timesteps and seconds. |
|  | Click frequency | 1.3 Hz | Click production rate, estimated from behavioral data. |
| | $v_{max}$ | $50^\circ / \Delta t$ | Upper bound for initial velocity. Randomly drawn at trial onset from uniform distribution $\mathcal{V}(0, v_{max})$. Falls within the range of rapid human head movements (Grossman et al., 1988). |
| | $\sigma_{base}$ | $0.5^\circ$ | Baseline perceptual measurement noise. Set to represent optimistic perceptual acuity. Finer than observed echolocation thresholds (Teng & Whitney, 2011; Teng et al., 2012; Thaler & Goodale, 2016), but allows the model's performance ceiling to be determined by reflectivity scaling rather than arbitrarily high baseline noise. |
| | Drift | $1^\circ$ | Motor variability parameter scaling movement control noise $\sigma_{motor}$. Set to produce realistic movement variability without overwhelming the control signal. |
| Theory-based | $P_0$ | $500^\circ{}^2$ | Uncertainty at trial onset. Large initial uncertainty reflects complete spatial ignorance at trial onset, consistent with Bayesian inference frameworks where priors are uninformative when no prior knowledge exists. |
| | $\hat{x}$ | $0^\circ$ | Initial KF estimate at trial onset. Updated after the first measurement. |
| | $Q$ | 0.1 | Process noise governing uncertainty change independent of measurements. Small nonzero value assumes static target while preventing KF overconfidence between measurements. |

### Experiment Parameters

In the simulated experiment, the participant begins facing forward (0^◦^ azimuth), and a target is placed randomly within the frontal plane between (± 90^◦^) at one of 500 pregenerated target azimuths. Each trial lasted up to 30 *s* or until the target was localized. Successful localization required meeting several criteria: (1) the head azimuth was within ±5^◦^ of the estimated target azimuth; (2) confidence estimate *P* plateaued at its asymptotic minimum value for the trial; and (3) the first two criteria were met for 3 consecutive measurements. Time was discretized at 10 Hz (equal to 300 time steps over 30 s) to match the update rate of the belief system.

The 500 pregenerated trial parameters were run on each of the three experimental conditions (Control, Big Target, Small Target). In the control condition, the model implemented the original experiment’s “No-Click” condition by preventing measurement updates to the KF. The Big and Small Target conditions differed only in the size of the target to localize. The model implemented this by scaling the simulated echo proportionally to the change in surface area — from a factor of 1 for Big Target to 0.04 for Small Target trials. To capture this difference in target dimensions, the model used a reflectivity scaling factor that modulated measurement quality. The rest of the parameters remained the same.

### Acoustic Measurement and Processing Module

#### Audio library

Using waveform and directivity models of real-world echolocation clicks (Thaler et al., 2017) convolved with head-related transfer functions (HRTFs) from a KEMAR acoustic manikin (Gardner & Martin, 1995), we generated a library of binaural acoustic echoes organized by angle with 1^◦^ resolution. In the test conditions, the model uses this library to simulate the acoustic echo corresponding to the angular error between the current head direction and the target. The echo ITD is computed using the ITD processor (see next subsection) to update the KF belief with a new target location estimate and the strength of the belief. The audio matching tolerance is set to 0.5^◦^ to ensure that azimuthal ITD estimates are rounded to the nearest degree.

#### ITD processor

The purpose of the ITD processor is to extract acoustic cues from binaural audio and convert them to spatial (azimuth) estimates. We extracted binaural cues by cross-correlating windowed segments of the left and right audio channels around the main echo peak. To convert the resulting ITD to azimuthal angle, we used a spherical head model assuming a head radius of 9 cm, an ear distance of 18 cm and the speed of sound (*c* = 343 *m/s*):

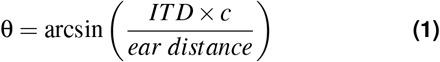

#### Signal quality

We calculated a reliability measure, or base quality, for each echo measurement, as the normalized strength of the cross-correlation:

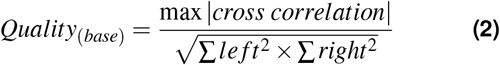

The range of base quality is between [0, 1], where 0 represents a noisy signal with no correlation and 1 represents a signal with high quality and a clear ITD peak. We then adjusted base quality for target reflectivity, with signal strength scaling with the square root of the reflecting area:

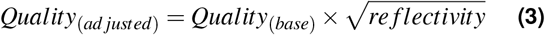

Measurement quality was affected through two mechanisms: First, small targets produced weaker echoes with worse cross-correlation quality (captured by *Quality*_(*adjusted*)_). Second, these weaker echoes increased perceptual noise in angle estimation. The standard deviation of the measurement noise scaled inversely with reflectivity:

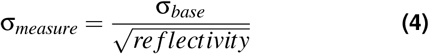

Thus, the small target produces noisier, less reliable measurements, which leads to slower convergence and lower successful localization rates.

### Belief Module

The KF maintains a probabilistic estimate of the target location and quantifies uncertainty in that estimate. This estimate directly affects the motor control system.

#### State representation

State variable 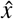 represents the current best guess of where the target is. At the start of the trial, 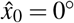. Uncertainty *P* (variance) quantifies confidence in the estimate, with lower values indicating higher confidence, leading to narrower exploratory head movements. At the start of the trial, *P*_0_ = 500^◦2^, signifying a large uncertainty.

#### Measurement model

When an echo is detected, the ITD processor extracts a noisy estimate of target azimuth *z*, modeled as:

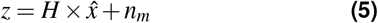

where *H* is the observation matrix. Here, *H* = [1], since we have a 1D model and we measure angle directly; *n*_*m*_ ∼ *N* (0, *R*_*k*_) represents measurement noise with variance *R*_*k*_. The KF then uses this measurement to update its belief about *x*. In this model, measurement noise *R*_*k*_ is adaptive, i.e., measurement reliability varies with the quality of the echo:

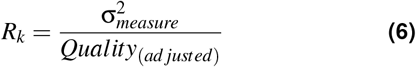

Good echoes result in low *R*_*k*_, which leads to the KF trusting the measurement more.

#### Process model

The target is assumed to be static, thus the process model is determined by the simple equation:

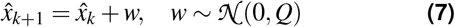

where 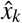 represents the estimated target azimuth at time step *k* and *w* is process noise. Although the target is static, small process noise *Q* prevents overconfidence during periods with no acoustic measurements. This allows uncertainty *P* to slowly grow over time.

#### KF algorithm

At each time step Δ*t* = 0.1*s*, the KF executes two stages:

1. **Prediction** (executed in all conditions):

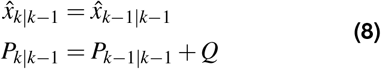

The predicted state 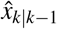 remains unchanged (static target), while the predicted uncertainty *P*_*k*_|_*k*_−_1_ increases slightly due to process noise *Q*.
2. **Measurement update**: This step only happens when there is a click made and an echo measurement is obtained as a consequence. When a click produces a valid echo measurement *z*_*k*_, the filter computes:

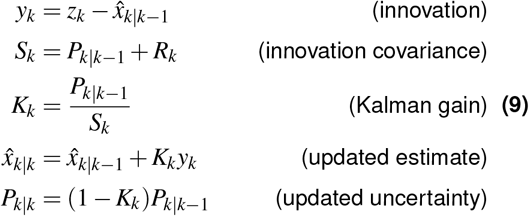

The observation matrix *H* = [1] simplifies these equations, since we observe target azimuth directly in this 1D model. The Kalman gain *K*_*k*_ determines how much the new measurement influences the estimate: values near 1 indicate high trust in the measurement (low *R*_*k*_ or high uncertainty *P*), while values near 0 indicate the prior estimate is retained (high *R*_*k*_ or low uncertainty *P*).

### Motor Control Module

The motor control system generates head movements through a combination of velocity and direction anchored to the KF estimate, with proportional gain and damping parameters, ensuring more realistic convergence dynamics.

#### State variables

The controller maintains three state variables: head azimuth θ (^◦^), angular velocity *v* (^◦^*/timestep*) and movement direction *d* ∈ {−1, +1}. At trial onset, direction is randomly initialized (*d* = ±1 with equal probability), and velocity is set to *v*_0_ = *d* × *V* (0, *v*_*max*_), where *v*_*max*_ = 50^◦^*/timestep*, i.e., the model randomly draws from a uniform distribution of velocities ranging from 0 to 50.

#### Desired target position

The controller determines a desired target position θ_*target*_ toward which the head should move. This desired position is determined by the KF current best estimate of target location 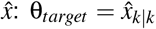. Before the first measurement happens, 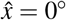, and the head starts exploring around that forward-facing direction. As measurements arrive at the next time steps and the belief is updated, θ_*target*_ shifts accordingly, causing the center of exploration to follow the estimated target location.

#### Movement dynamics

Rather than updating the head position directly using errors between current estimate θ and desired position θ_*target*_, the controller modulates velocity through a proportional control approach with adaptive damping. Thus, velocity *v* serves as an intermediate state variable representing the velocity of the head. At each timestep, the position error is computed as:

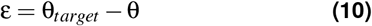

Velocity is then updated using proportional gain *K*_*p*_ and damped to prevent overshooting:

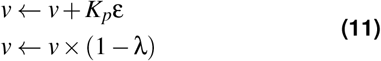

where λ is the damping coefficient. The proportional gain and damping adapt based on whether echo feedback is available:

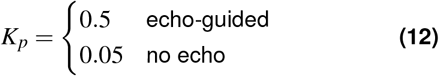

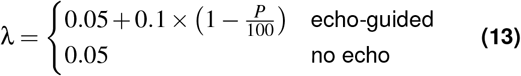

In echo-guided mode, damping increases as uncertainty *P* decreases, resulting in more precise, responsive movements when confident and smoother, broader exploratory behaviors when less confident. Velocity is then clipped to physical linits (|*v*| ≤ *v*_*max*_) and the head position θ is updated based on the new velocity:

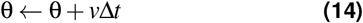

This approach with velocity as an intermediate state variable introduces momentum, lag, and more biologically plausible overshoot-correction behaviors rather than instantaneous position tracking.

#### Boundary handling

If head position reaches the physical rotation limits (|θ| ≥ 90^◦^), two mechanisms are triggered to ensure smoother reversals. First, position is clipped to the boundary:

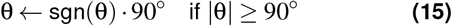

Second, velocity is reversed and damped by 30% to prevent elastic-like bouncing:

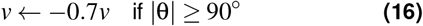

#### Movement control noise

To simulate natural motor control variability, small random perturbations are added to the final head position at each timestep:

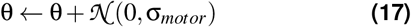

where σ_*motor*_ represents movement control noise, scaled by the drift parameter.

This control architecture produces movement patterns qualitatively similar to human echolocators, where broader exploratory sweeps happen early in trials when uncertainty is high, gradually narrowing down to smaller, more targeted movements as the target location is refined.

## Analysis

To assess model performance across conditions, we evaluated the following behavioral outcomes: target localization performance, individual head trajectories over time, localization accuracy (distribution of final errors) and convergence dynamics, and performed statistical analyses to compare performance between conditions.

### Target localization performance

The model counted a target as ‘found’ when (1) confidence P declined by < 5% for 3 consecutive measurements, and (2) the head was within 5^◦^ of estimated target azimuth. This represented ‘subjective’ detection - when it believed it had successfully localized the target. For an objective evaluation, we also computed the proportion of trials in which the final head position was within 5^◦^ of the actual target. For trials where the model declared detection, we calculated response time (RT) as trial duration from onset to the moment of subjective detection.

### Individual head trajectories

To visualize search and convergence behaviors, we plotted head orientation over time for individual trials, time-locked to trial end. Trajectories were plotted relative to target position to reveal patterns independent of absolute target location.

### Localization accuracy (distribution of final errors)

We quantified final localization accuracy using the error at the final timestep of the trial. For each condition, we computed mean, median, and standard deviation of final errors across all trials. Accuracy distributions were visualized as histograms to reveal whether errors clustered near zero or showed broader distributions. We compared final errors across conditions using independent samples t-tests, and tested for differences in precision (variability) using Levene’s test for equality of variances. The variance ratio between the Small and Big target conditions quantified the relative precision between conditions.

### Convergence dynamics

We operationalized convergence efficiency as error reduction over the course of a trial, yielding a normalized measure of convergence speed. We then visualized convergence dynamics by plotting mean absolute error (MAE) as a function of time from trial start. To handle varying trial durations, we carried forward each trial’s final error value for timepoints after detection, allowing computation of averages across all trials per condition.

### Statistical analysis

We compared target localization rates across conditions using chi-square tests. For continuous metrics (RT, final error, and convergence rate), we used independent samples t-tests and quantified effect sizes using Cohen’s *d*. For localization precision (SD of final errors), we used Levene’s test for equality of variances. We conducted pairwise comparisons between the conditions, focusing primarily on the differences between the test conditions (Big Target, Small Target).

## Results

### Target localization performance

#### Objective localization rates

The rates of objective target localization for the three conditions were as follows: 0% for the Control condition omitting clicks, 81.4% for the Big Target condition (407/500 trials) and 38.4% for the Small Target condition (192/500 trials) (Fig. 4a). Chi-square tests revealed a main effect of target size (Control vs Big Target: χ^2^ = 581.361, *p* < .001; Control vs Small Target: χ^2^ = 154.831, *p* < .001; Big Target vs Small Target: χ^2^ = 190.659, *p* < .001)

**Fig. 4.**
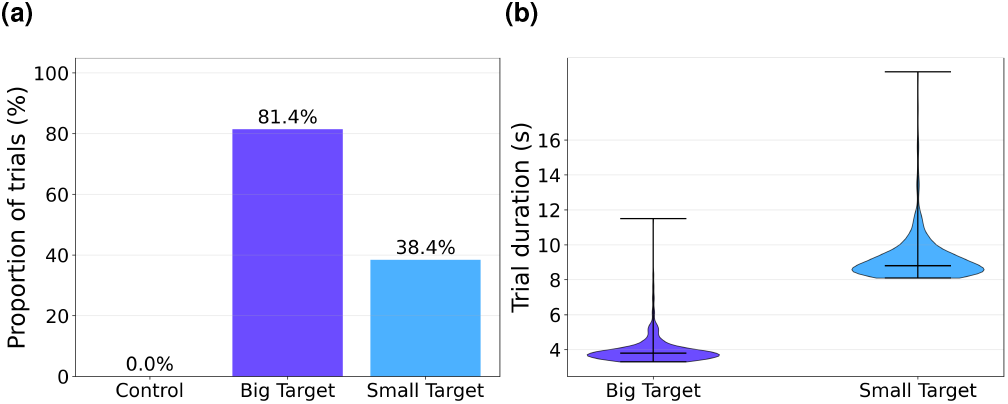
Target localization performance: localization rates (a) to within 5◦ of target; and RT (b) for target localization across test conditions. Violin plots show the distribution of response times (trial duration from onset to subjective detection) for Big Target (left, dark blue, n=500) and Small Target (right, light blue, n=500) conditions. Violin width represents probability density at each response time value. Internal black lines indicate median (horizontal line) and range (vertical lines extending to minimum and maximum values).

#### Response times

Response times were significantly shorter in the big target condition (*M*_*big*_ = 3.95*s, SEM*_*big*_ = 0.03*s*) than in the small target condition (*M*_*small*_ = 9.21*s, SEM*_*small*_ = 0.06*s*), *t*(998) = −76.880, *p* < .001, *d* = −4.867) (Fig. 4b)

#### Individual head trajectories

Figure 5 shows angular trajectories for each condition, with 50 trials randomly subselected for clarity. Trajectories show head azimuth relative to target position (head orientation minus target azimuth), with the red dashed line at 0°indicating perfect alignment to target. This representation reveals approach patterns independent of absolute target location. In the Control condition, trajectories show a lack of systematic convergence in the absence of measurements. In contrast, both Big and Small Target conditions show directed convergence toward the target, though Small Target trials exhibit greater variability and wider oscillations during approach.

**Fig. 5.**
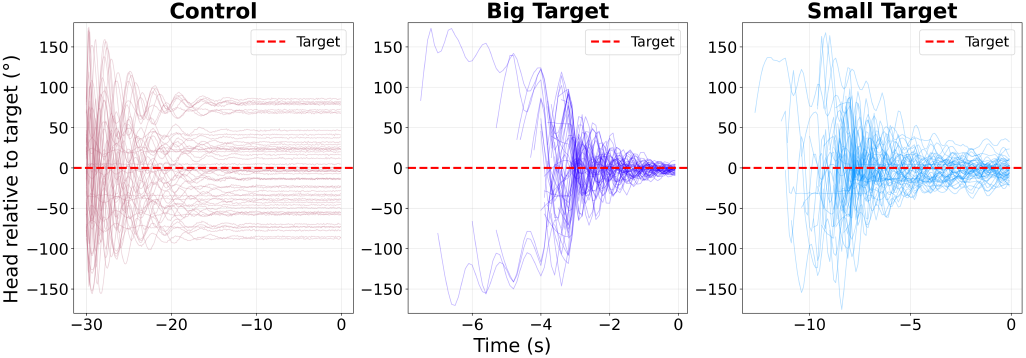
Individual trial trajectories across conditions. Head orientation relative to target (0°= alignment, dashed red line) for 50 trials per condition, aligned to trial end (t=0). Control (left, red) shows no convergence; Big Target (middle, dark blue) shows rapid convergence; Small Target (right, light blue) shows slower convergence with greater variability.

#### Localization accuracy

The analysis of final error distributions revealed significant differences in localization precision across conditions (Fig. 6). In the Control condition, final errors were uniformly distributed across the full angular range (*M*_*ctrl*_ = 46.7^◦^ ± 26.4^◦^), indicating no systematic convergence on the target. In contrast, the Big Target condition produced highly accurate localization patterns with final errors tightly clustered around the target (*M*_*big*_ = 3.5^◦^ ± 8.1^◦^). The Small Target condition showed lower accuracy with a broader error distribution (*M*_*small*_ = 9.6^◦^ ± 14.4^◦^). Small Target localization was both less accurate (*t*(998) = −8.280, *p* < .0001, *d* = −0.524) and less precise (Levene’s *W* = 38.040, *p* < .0001, Variance ratio (Small/Big) = 3.18) than localization in the Big Target condition.

**Fig. 6.**
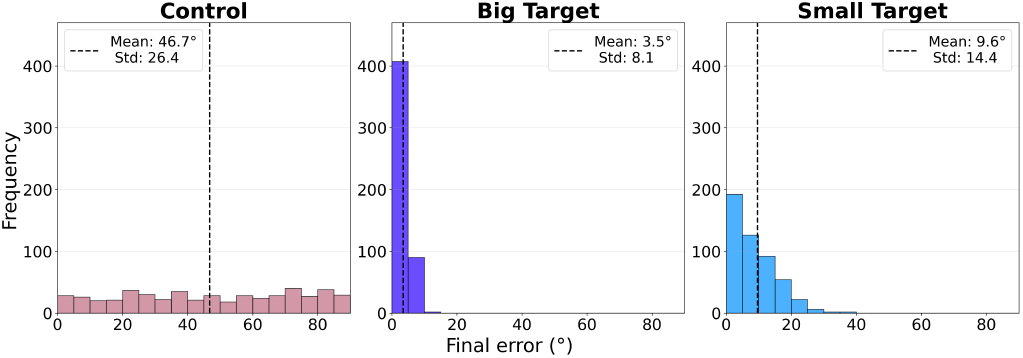
Final error distributions and variance differences between Control (left, red), Big Target (middle, dark blue) and Small Target (right, light blue) conditions. The vertical dashed black line represents the mean of the distribution.

#### Convergence dynamics

Analysis of error reduction over time revealed convergence patterns across conditions (Fig. 7). In the Control condition, MAE remained stable at approximately 45^◦^ throughout the trial, confirming the absence of convergence without acoustic feedback. Both Big and Small Target conditions showed systematic error reduction, but with different rates. Big Target trials demonstrated more rapid convergence compared to Small Target trials, with error reduction to within 5^◦^ at 11.02^◦^*/*s vs. 4.05^◦^*/*s, respectively (*t*(998) = 20.249, *p* < .0001, *d* = 1.282). This shows that target size affects not only the accuracy of localization, but the efficiency of the localization process in general.

**Fig. 7.**
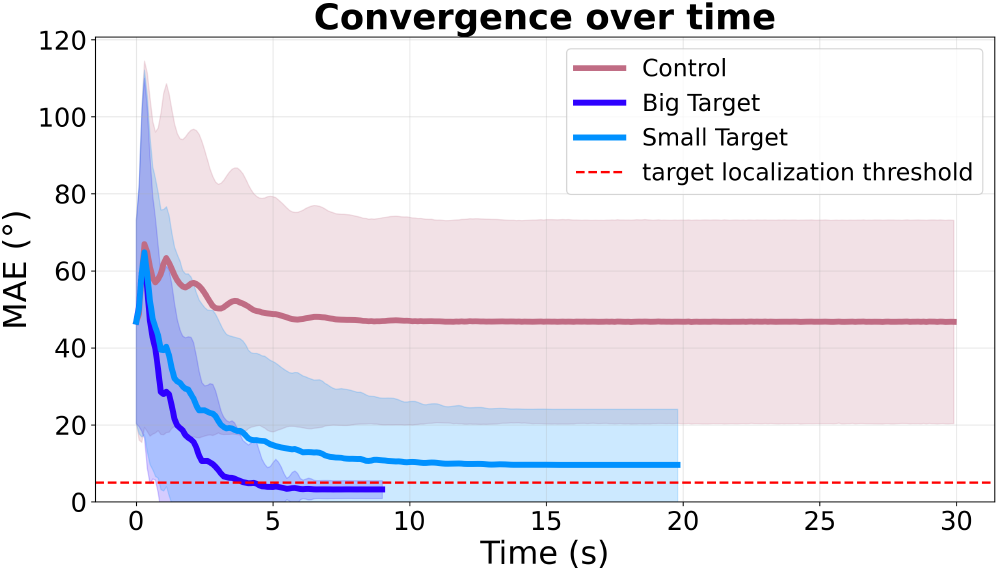
Convergence (error reduction over time). Reductions in MAE over time for Control (red), Big Target (dark blue) and Small Target (light blue) conditions. Solid lines show mean error; shared regions represent ±1 SD; the horizontal red dashed line represents the spatial threshold set to 5◦ of the target. MAE: Mean Absolute Error.

## Discussion

Our model of echolocation based on a KF successfully localizes targets using echoacoustic feedback. Comparisons of model perfomance across three conditions (Control with no echoaoustic feedback, Big Target and Small Target with echoacoustic feedback) demonstrated that localization performance and dynamics depend on target size manipulation and on echo measurement availability. The most interesting key effects were based on the comparison of the two target size conditions (Big vs. Small): localization rates (81.4% vs. 38.4%), response times (3.95 s vs. 9.21 s), response precision (8.1^◦^ vs. 14.5^◦^), final errors (3.5^◦^ vs 9.6^◦^) and convergence rates (11^◦^*/*s vs 4^◦^*/*s) demonstrate improved performance in the Big Target condition compared to the Small Target condition, implying that target reflectivity played a critical role in performance.

The model reproduces the target dependence of localization performance observed in a human expert echolocator, with reliably faster, more accurate, and more precise localization behavior for Big vs. Small targets. Localization precision and trial length qualitatively approximates realistic echolocation behavior in the original paradigm (Patel et al., 2024). However, model performance is quicker and more precise than human performance: *SD*_*human*_ = 13.6^◦^ vs *SD*_*model*_ = 8.1^◦^ for the Big Target condition, *SD*_*human*_ = 32.3^◦^ vs *SD*_*model*_ = 14.5^◦^ for the Small Target condition, and *SD*_*human*_ = 45.9^◦^ vs *SD*_*model*_ = 26.4^◦^ in the No-Click condition (Patel et al., 2024). Trial lengths are also significantly higher in the Small Target condition compared to the Big Target condition (3.9*s* vs 9.2*s*, respectively), consistent with human performance (9.3*s* vs 23.2*s* on Big and Small trials, respectively, 13.9*s* difference; *p* < 0.001) (Teng et al., 2026).

Although comparable to human behaviors, model performance surpasses that of human echolocators, as is often the case with ideal observers(Kao et al., 2004). The KF may lack integration strategies that humans might employ when managing noisy sensory information. Humans face biomechanical constraints, motor variability, and cognitive processing loads that may reduce efficiency, while the model assumes perfect sensorimotor coordination: motor commands execute without delay or error, and clicks are produced at fixed regular intervals. Unlike human participants, the model does not get mentally exhausted and its tongue does not tire. An approach where the model is trained on real data may potentially resolve these constraints, as might more realistic modeling of sensory and motor noise (Yang et al., 2016). Future work should consider data-driven approaches to parameter estimation, where model parameters are fitted to empirical human echolocation trajectories rather than selected a priori. Exploring hierarchical frameworks such as recurrent neural networks (Barak, 2017; Bolhasani et al., 2025; Durstewitz et al., 2023) may be more fitting for human sensorimotor behavior. Despite these limitations, the KF-based model successfully captures the qualitative pattern of uncertainty-driven exploration, demonstrating that a Bayesian-based inference approach coupled with adaptive motor control provides a plausible mechanistic account of human echolocation behavior. This framework may thus be used to generate simulated behavioral data to train and test future data-driven models before (or in parallel with) time-consuming collection of real human data.

## Conclusion

The presented echolocation model couples a KF with a motor controller and captures key features of human echolocation behavior, with target surface area affecting performance via measurement noise. A 25-fold reflectivity difference between the Big and Small Target conditions produced 43% lower localization rates, 133% longer response times and 3.2-fold increases in error variance for the Small Target. These features of the simulated performance, and the sinusoidal pattern of echo-guided head movements, qualitatively reproduce previously observed human behavior. These effects emerged naturally from the model’s probabilistic sensory integration without the need for explicit strategy changes. Importantly, these dynamics were produced with very few explicit links to observed human data. The model generates testable predictions about the relationship between target dimensions and localization dynamics. More broadly, our framework demonstrates that a KF coupled with uncertainty-driven motor control provides a computationally plausible and biologically motivated account of active sensorimotor behavior, and therefore may be used as a well characterized starting point from which to implement more detailed simulations. Future work exploring other computational frameworks and direct validation against human behavioral and neural data will further strengthen the connection between computational models and human active sensing strategies.

